## Supplementary Figures and Tables for "A subset of viruses thrives following microbial resuscitation during rewetting of a seasonally dry California grassland soil"

| Metagenome type | Method | Presumed composition | Quantity |
|---|---|---|---|
| Unfractionated metagenome | Direct DNA extraction from soil (Figure 1A) | Bacterial, archaeal, eukaryotic, and viral sequences from MAGs and vOTUs (which may be intracellular or extracellularized). | 24 |
| SIP-fractionated metagenome | DNA extracted from soil and ultracentrifuged in a CsCl gradient to capture distinct DNA-density fractions for quantitative stable isotope probing (qSIP) (Figure 1B) | $H_2^{18}O$-incorporating MAG and vOTU genomes in heavier DNA fractions i.e., considered active. | 210 |
| Virome | Small-fraction metagenome, i.e. viral-enriched sequences: soil buffered, 0.2 µm filtered, concentrated, treated with DNase. DNA extracted from 0.2 µm effluent (Figure 1A) | Sequences from < 0.2 µm membrane-bound or encapsulated genomes i.e. Viral-like particles (VLPs) or virions; and ultrasmall cells such as CPR bacteria and nanoarchaea | 18 |

**Supplementary Table 1 | Mixed ‘omic methods enable detection of genomes and distinguishing as abundant, active, and virions.**

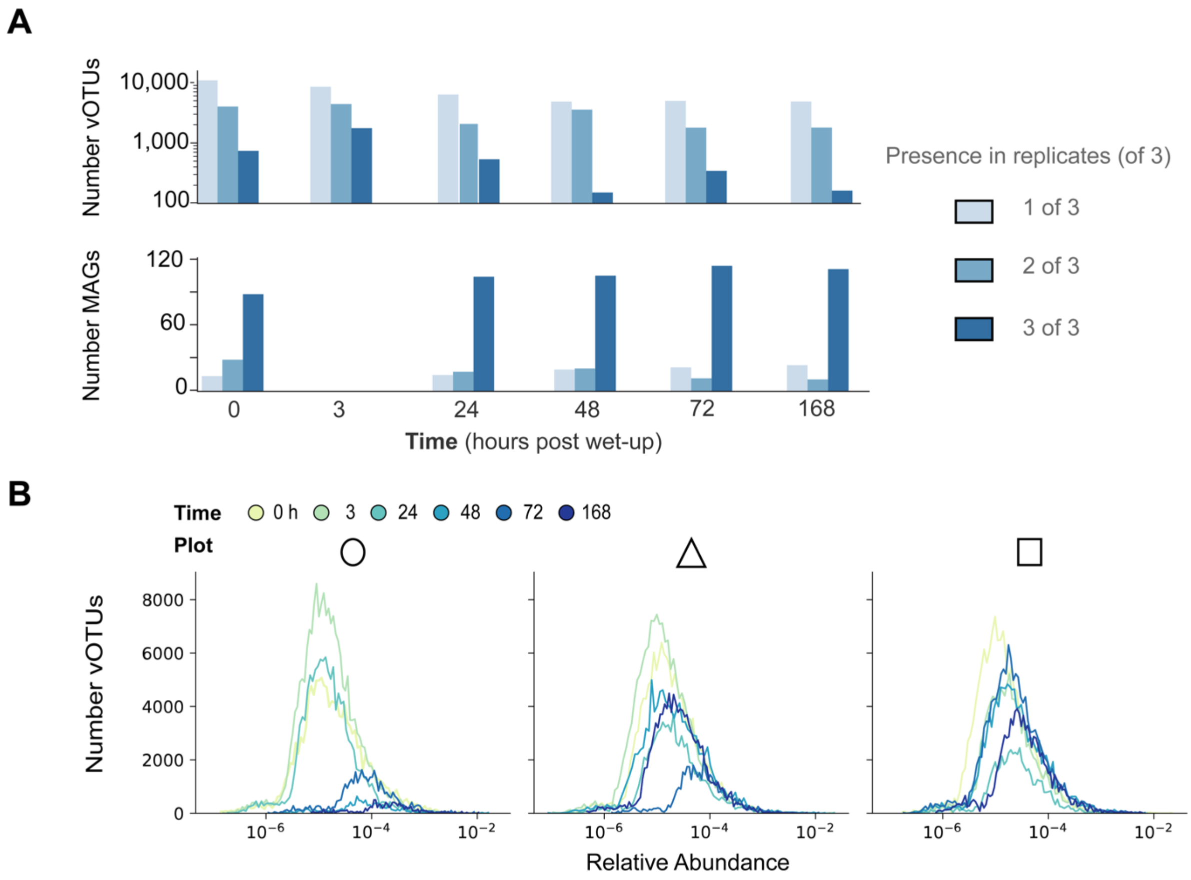


**Supplementary Figure 1 | Viral heterogeneity through space.** (A) Barplots visualize the number of (top) recovered vOTUs (log scale) or (bottom) MAGS found in one (lightest blue), two, or three (darkest blue) of three replicate microcosms. (B) Distribution of vOTU relative abundances per time point (colored curve) graphed by field plot (signified by circle, triangle, and square used throughout). The y axis represents the number of unique vOTUs detected per time point.

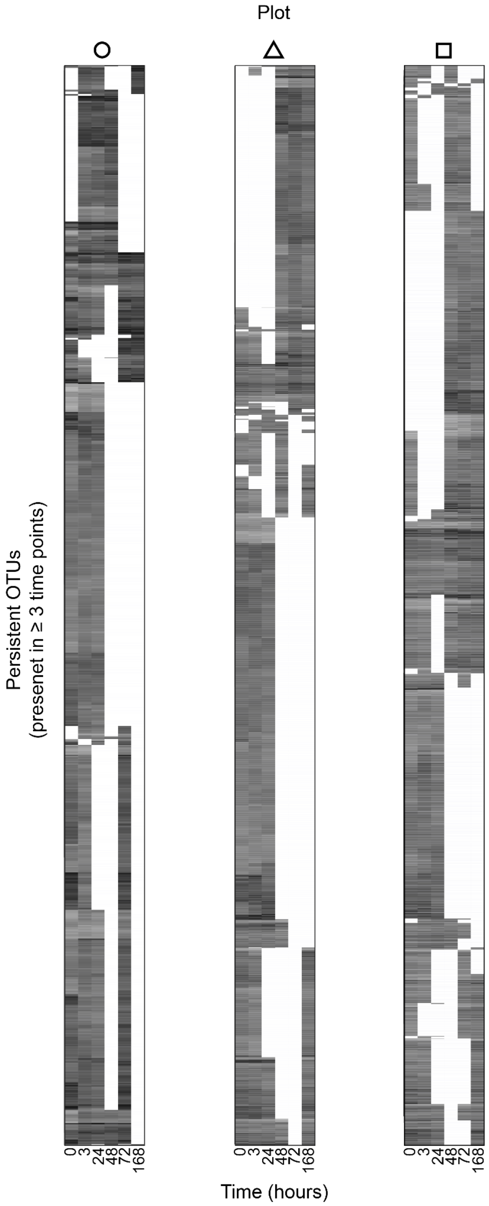


**Supplementary Figure 2 | Temporal changes in relative abundance of persistent viral populations (vOTUs).** Each heatmap corresponds to one of the three field plots sampled and is hierarchically clustered according to vOTU relative abundance. Time is shown on the x-axis of each plot (columns); Each row (y-axis) corresponds to a unique vOTU and its relative abundance through time. All vOTUs were present in each given plot in at least three time points, i.e., "persistent" vOTUs. Circles, triangles, and squares correspond to the field plots in the PCoA legend (Figure 2).

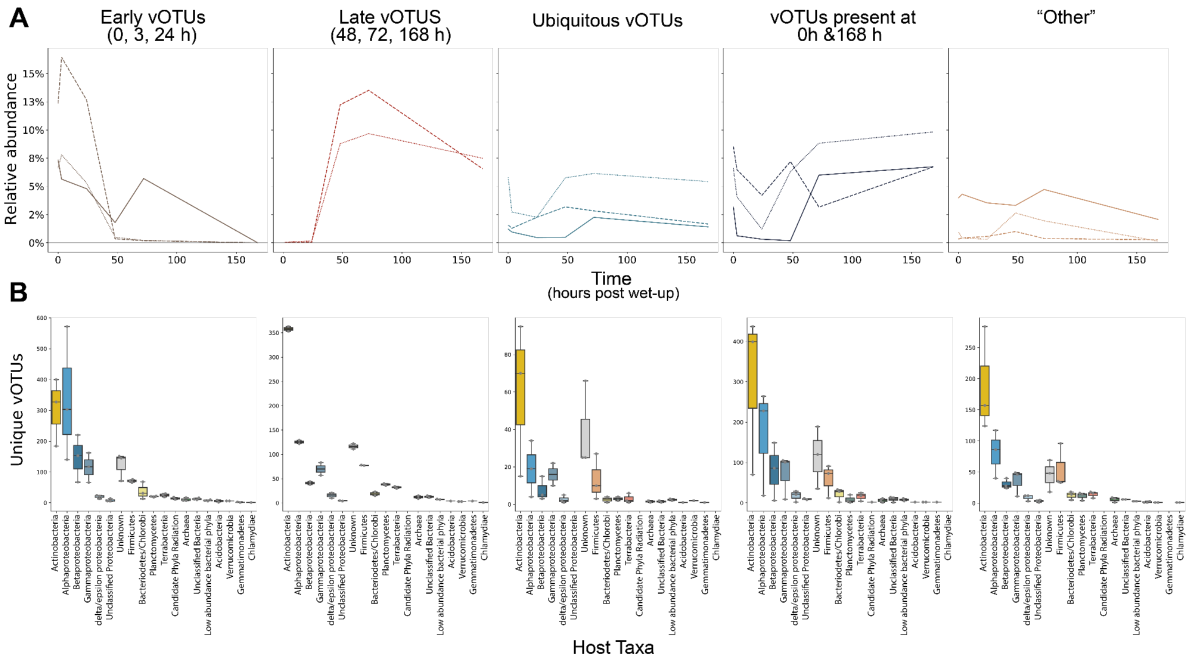


**Supplementary Figure 3 | Viral dynamics following soil wet-up classified by temporal response. (A)** Each plot shows the aggregated relative abundance per plot of the persistent vOTUs that respond early (0, 3, 24h); late (48, 72, 168h); ubiquitously (at all time points); present at 0h, 168h, and one other time point; and other. Relative abundance is shown as the percent of total reads per virome. **(B)** Boxplots graphed per response category showing the range of vOTU counts (y axis) separated and colored by predicted host taxonomic groups (x axis) across all three plots. The x axis is sorted by rank, but with all Proteobacteria groups adjacent to one another. The box plots represent 75% of the data with the median as a line, and whiskers represent 90% of the data.

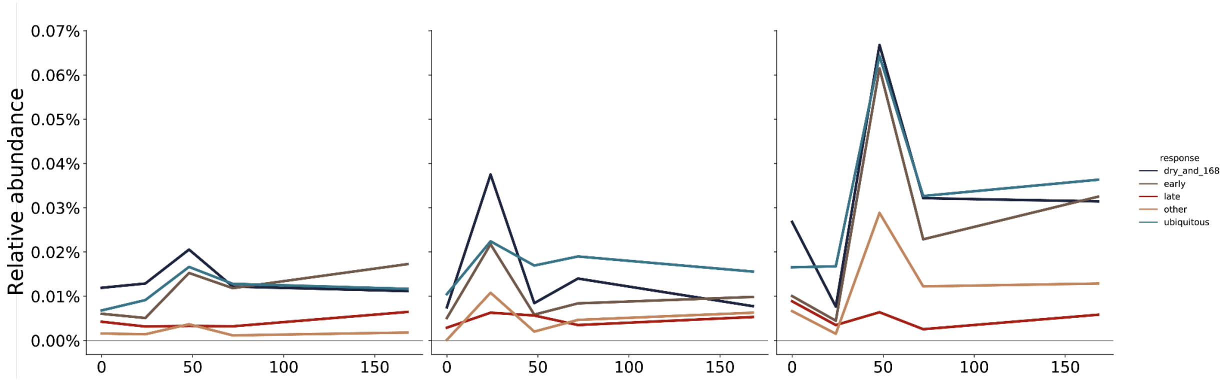


**Supplementary Figure 4 | Field plot-specific viral responses in unfractionated metagenomes.** Each graph represents unfractionated metagenome reads from one field plot mapped to the set of viruses of a specific response category defined in the virome (Sup. Fig. 3). Relative abundance represents the percent of total aggregated reads mapped to vOTUs of a given category.

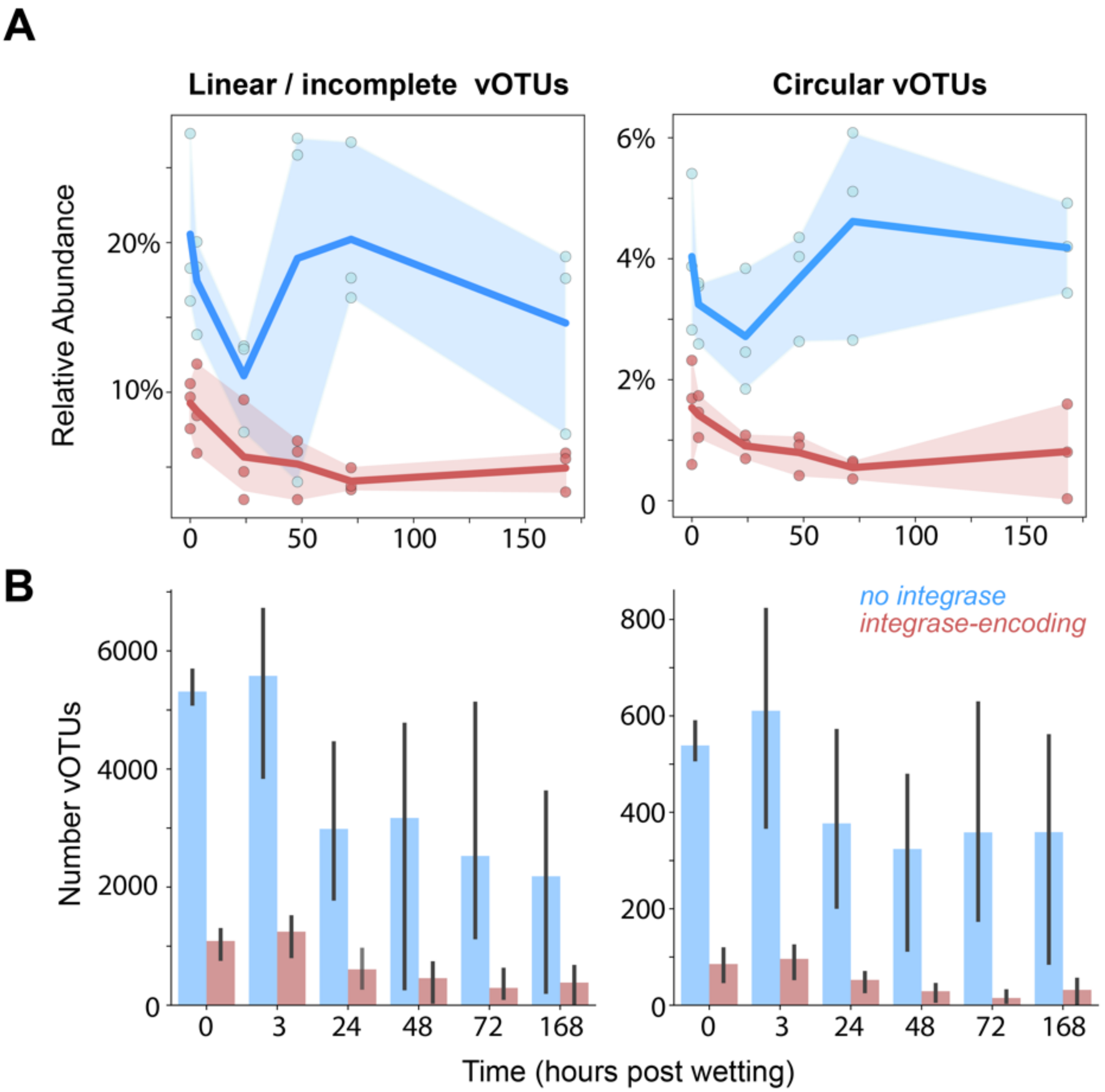


**Supplementary Figure 5 | Relative abundance and richness of integrase-encoding vOTUs through time following soil wet-up.** vOTUs were split by whether they encode an integrase gene (red) or not (blue) and whether the vOTU is predicted to circularize (right set of graphs) or is linear and perhaps fragmented (left set of graphs). (A) Aggregated relative abundance (percent of total sample reads) of integrase-encoding or not integrase-encoding vOTUs. (B) Counts of integrase-containing and non-integrase containing vOTUs.

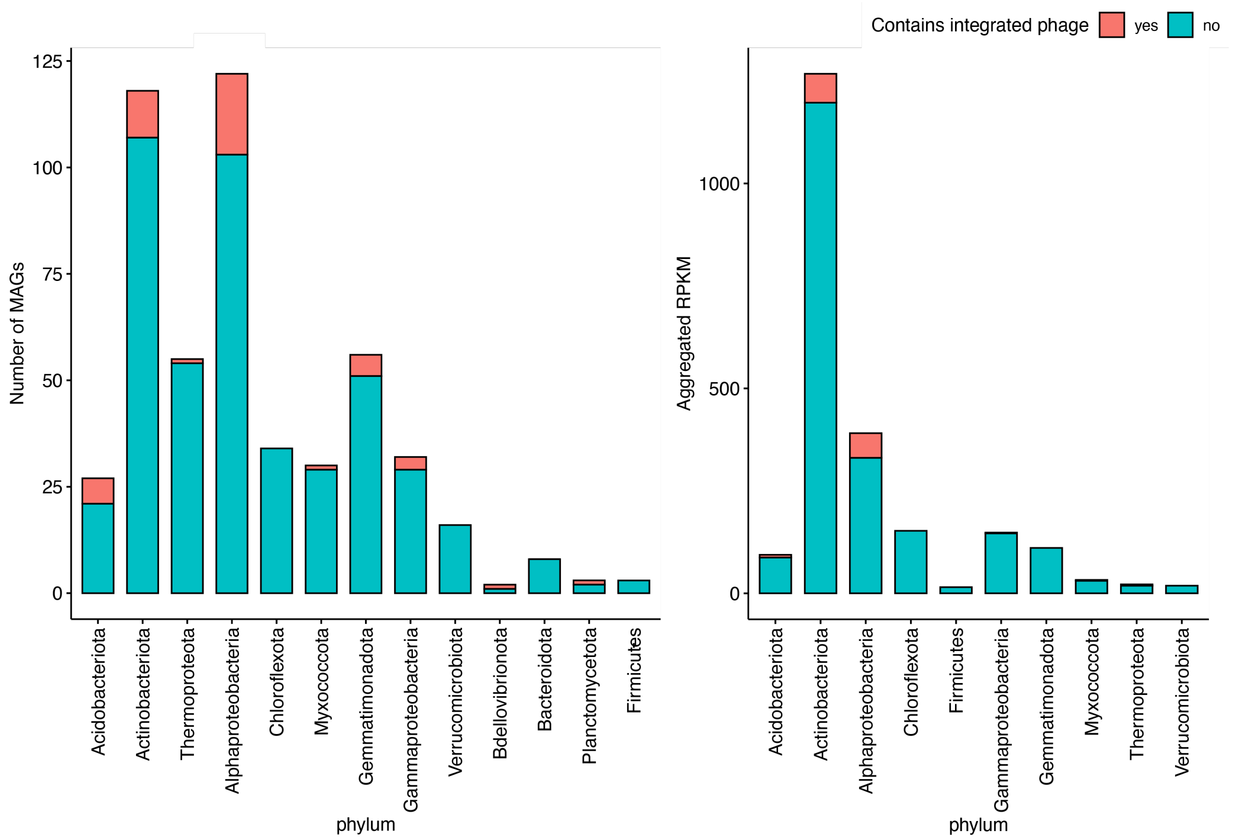


**Supplementary Figure 6 | Prophage detected in bacterial genomes (MAGs).** For each MAG of a given phylum the number of MAGs (left) and the relative abundance in RPKM of those MAGs is shown according to whether the MAG contains an integrated phage (red) or not (blue).
